# CASM regulates p62/KEAP1/NRF2 antioxidant responses to lysosome damage

**DOI:** 10.64898/2026.07.08.736046

**Authors:** Yasmeen Safayd, Karen E Anderson, Sophie Blagg, Joanne Durgan, Ruchi Sharma, Oliver Florey

**Affiliations:** Signalling Program, Babraham Institute, Cambridge, CB22 3AT, UK; Stemnovate LTD, Babraham Research Campus, Cambridge, CB223AT, UK

## Abstract

Effective lysosome function is essential for health, and declines with ageing and disease. Upon lysosome damage, cells mount a complex stress response to restore homeostasis. Membrane ATG8ylation plays a central role, orchestrating lysosome repair, replacement and removal, either directly at the damaged membrane, via CASM (<u>c</u>onjugation of <u>A</u>TG8s to <u>s</u>ingle <u>m</u>embranes), or at nascent autophagosomes, during lysophagy. Here, we identify a novel role for CASM in driving an antioxidant response to lysosome stress. Following damage, CASM regulates both lysosome ubiquitination and p62 recruitment. Membrane-associated p62 undergoes S349-phosphorylation, which permits KEAP1 sequestration, thereby releasing master transcription factor, NRF2. Liberated NRF2 translocates to the nucleus and promotes transcription of antioxidant and detoxifying genes, co-ordinating a cytoprotective response, in a CASM-dependent manner. These findings position CASM as a major upstream response to lysosome stress and uncover the p62/KEAP1/NRF2 axis as a novel effector pathway, termed SOLAR (SQSTM1/p62 Oligomer-mediated lysosome antioxidant response).

**Highlights:**

- CASM recruits p62/SQSTM1 as an effector protein during lysosome damage
- CASM acts upstream to enhance ubiquitination of damaged lysosomes
- CASM-recruited p62 engages KEAP1/NRF2 to drive antioxidant gene transcription (SOLAR)

## Introduction

Lysosomes are membrane-bound, acidic organelles with multiple fundamental cellular functions. They represent the primary degradative compartment of the cell, mediating the breakdown and recycling of intracellular and extracellular materials ^1^. They also serve as vital signalling hubs, that regulate nutrient sensing, metabolism, and cellular stress responses ^2^. As such, maintenance of lysosomal integrity is essential for cellular homeostasis and survival.

Lysosomal damage can be induced by diverse stressors, including: lysosomotropic agents, crystalline and aggregated materials, membrane-permeabilising toxins, pathogen infection, genetic diseases and ageing ^3,4^. Disruption of lysosomal membrane integrity releases hydrolytic enzymes, iron and other acidic luminal contents into the cytosol, promoting inflammation, cell death signalling, and increased reactive oxygen species (ROS) production, that drives oxidative damage and cellular dysfunction.

In response to lysosome stress and damage, cells harness a variety of responses to restore homeostasis, including; lysosome membrane repair ^5–8^, biogenesis of new lysosomes ^9,10^ and clearance of damage through lysophagy, a form of selective autophagy ^11,12^. Lysophagy is initiated following lysosomal membrane damage, which exposes luminal glycans to the cytosol, triggering recruitment of damage sensors such as galectins ^13^. Damaged lysosomes become decorated with ubiquitin chains, through the action of specific E3 ubiquitin ligases ^14-^18, creating a platform for the recruitment of selective autophagy receptors (SARs), including p62, NDP52, NBR1, TAX1BP1 and OPTN ^19,20^. These cargo receptors bind to both ubiquitin, through ubiquitin-associated domains (UBA), and ATG8 family proteins, through LC3-interacting regions (LIRs), thereby bridging the damaged lysosomes to the autophagy machinery. In parallel, autophagy initiation complexes are recruited to promote phagophore nucleation and expansion around the damaged organelle ^11,21^. Conjugation of ATG8 proteins to lipids on the growing autophagosome, via membrane ATG8ylation, facilitates cargo capture and autophagosome maturation, ultimately enabling delivery of damaged lysosomes to functional counterparts for degradation.

CASM (<u>c</u>onjugation of <u>A</u>TG8 to <u>s</u>ingle <u>m</u>embranes) is a parallel, autophagy-related pathway which is also closely linked to lysosome homeostasis ^22,23^. Unlike canonical autophagy (and lysophagy), which target ATG8 proteins to double-membrane autophagosomes, CASM promotes ATG8 lipidation directly onto the surface of endolysosomal membranes, mediated by the V-ATPase–ATG16L1 (VAIL) ^24–26^ and TECPR1-Sphingomyelin (STIL) axes ^27–29^. CASM appears to confer an early, rapid response to lysosomal stress, detecting loss of proton gradients (via VAIL) and membrane lipid changes (via STIL), and then coordinating a collection of downstream pathways involved in membrane repair, signalling, and lysosome quality control ^23^. These functions often involve engagement of ATG8 effector proteins, such as ATG2 ^30^, ALIX ^31–33^, FNIP1/2 ^34^, LRRK2 ^35^ and BLTP3A ^36^, but the mechanisms underlying the CASM-mediated lysosome stress response remain to be fully understood.

In the present study, we sought to investigate the mechanistic links between CASM, ATG8 lipidation and lysosome homeostasis. We reveal an unappreciated role for CASM in regulating the central p62 and ubiquitin responses to damaged lysosomes, and in co-ordinating adaptive antioxidant responses via KEAP1/NRF2-dependent transcription.

## Results

### CASM regulates p62 recruitment during lysosome damage

We hypothesized that CASM may modulate the lysosome damage response via direct recruitment of ATG8-effector proteins. To investigate a possible relationship between CASM and the selective autophagy receptors (SARs), p62 localisation was assessed in wild-type (WT), macroautophagy-deficient (ATG13 KO) or macroautophagy/CASM-deficient (ATG5 KO) cells, expressing GFP-ATG8 (LC3A), upon lysosome damage (LLOMe) (Fig. 1A, B). In WT and ATG13 KO cells, LLOMe induced robust GFP-LC3A puncta formation, indicating the acute ATG8 response to lysosome damage is largely independent of macroautophagy, consistent with recent reports ^30^. In parallel, p62 puncta were also induced, showing clear colocalisation with GFP-LC3A. In contrast, ATG5 KO cells exhibited a complete absence of LLOMe-induced GFP-LC3A puncta, as would be expected, and strikingly, a clear reduction in p62 puncta too. This observation suggests that counter to current models of lysophagy, p62 recruitment to damaged lysosomes requires ATG8 lipidation, and moreover, that this is independent of ATG13, consistent with a role for CASM.

**Figure 1.**
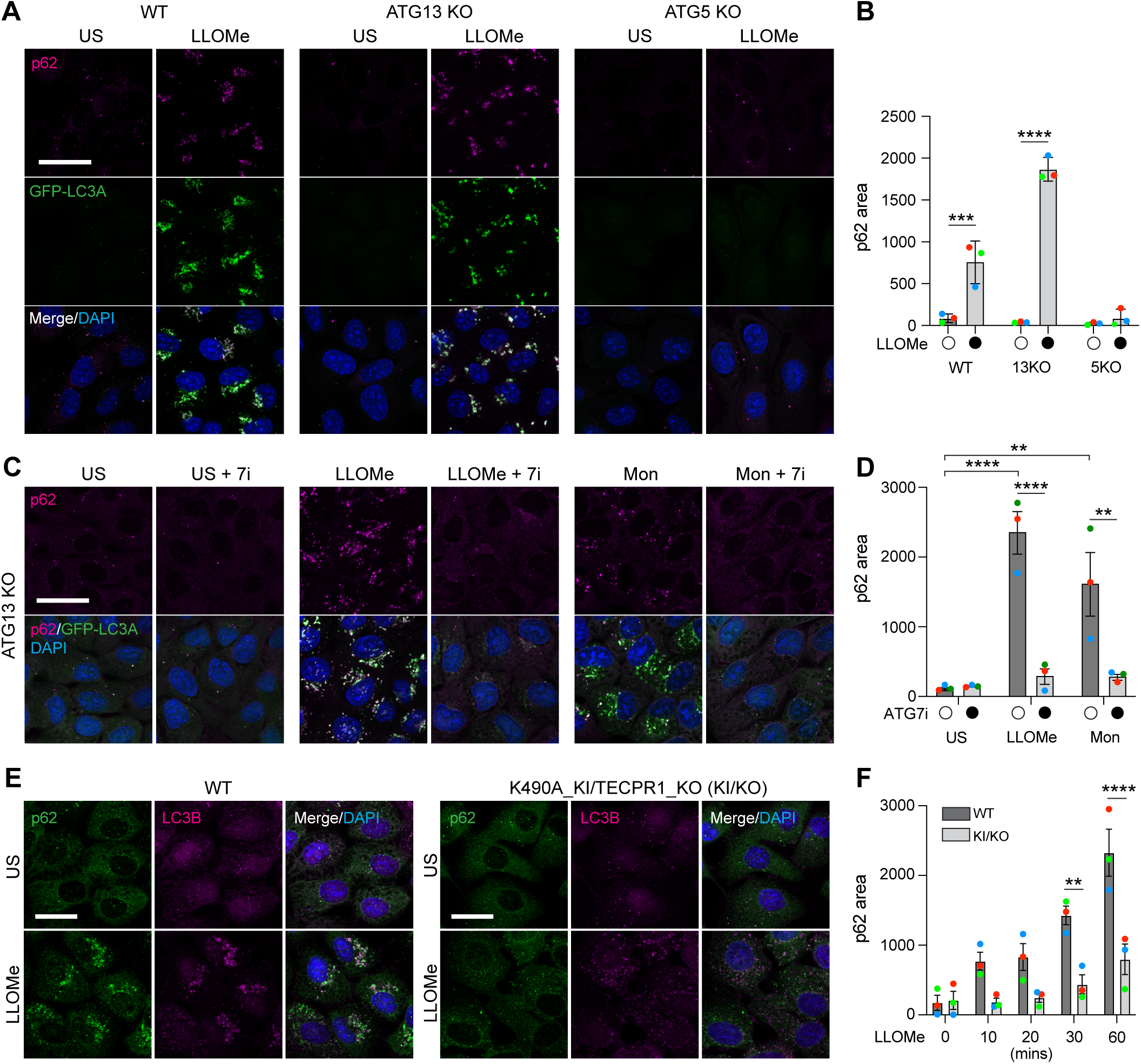
Lysosome damage induces CASM-dependent p62 recruitment. (A) Confocal images of p62 and GFP-LC3A in wild type, ATG13 KO and ATG5 KO MCF10A cells treated +/- LLOMe (250 μM, 30 mins). Scale bar: 25 μm. (B) Analysis of p62 puncta area from images in (A). Data represent means ± SEM from 3 independent experiments. *** p 0.0001, **** p<0.0001, 2 way anova with sidaks multiple comparison. (C) Confocal images of p62 and GFP-LC3A in ATG13 KO MCF10A cells treated with LLOMe (250 μM, 30mins) or monensin (Mon, 100 μM, 45 min) +/- pretreatment with ATG7 inhibitor (7i, 10 μM, 2 hr). Scale bar: 25 μm. (D) Analysis of p62 puncta area from images in (C). Data represent means ± SEM from 3 independent experiments. **** p<0.0001, ** p<0.0017, 2 way anova with sidaks multiple comparison. (E) Confocal images of p62 and LC3 in wild type and K490A TECPR1 KO (KI/KO) MCF10A cells treated +/- LLOMe (250 μM) for the indicated times. Scale bar: 20 μm. (F) Analysis of p62 puncta area from images in (E). Data represent means ± SEM from 3 independent experiments. *** p<0.0001, ** p<0.0026, 2 way anova.

To test this further, we employed an ATG7 inhibitor (ATG7i) as an alternative, pharmacological method to impede ATG8 lipidation; when used on ATG13 KO cells, ATG7i is effectively blocking CASM. As expected, ATG7i blocked GFP-LC3A puncta formation upon both lysosome damage (LLOMe) and lysosome stress (monensin) (Fig. 1C, D). Importantly, it also abrogated p62 puncta formation, consistent with the ATG5 KO. Thus, recruitment of p62 to damaged lysosomes bears the mechanistic hallmarks of CASM, showing dependence on ATG8-lipidation via ATG5/ATG7, but independence from ATG13.

To establish the CASM-dependence of p62 recruitment more directly, we engineered a CASM-deficient cell line (KI/KO), harbouring an ATG16L1 K490A knock-in (KI) mutation, combined with a TECPR1 KO, disrupting the VAIL and STIL axes of activation respectively (Fig. S1A-B). Importantly, while this cell line is unable to activate CASM, it retains competence for macroautophagy, responding to mTOR inhibition (PP242) in a Vps34-dependent manner (Fig. S1C, D), and forming WIPI2-positive puncta (Fig. S1E). Strikingly, p62 puncta formation was markedly reduced over a timecourse of LLOMe-treatment in KI/KO cells, compared to WT (Fig. 1E, F). Collectively, these findings provide evidence for a novel, upstream role for CASM in mediating p62 recruitment to stressed and damaged lysosomes.

### CASM-dependent recruitment is SAR-selective and context-dependent

To determine whether CASM engages p62 specifically, or SARs more broadly, we next examined additional family members, NBR1 and NDP52. Interestingly, NBR1 recruitment exhibited similar behaviour to p62, albeit at lower levels, being independent of macroautophagy, but requiring ATG8 lipidation (Fig. S2A, B). In contrast, LLOMe-induced NDP52 puncta formation remained robust in ATG13 KO cells, and occurred independently of ATG8 lipidation (Fig. S2C, D). These findings indicate that SAR recruitment is differentially regulated during lysosome damage, with some family members dependent on CASM, and others independent of any ATG8ylation.

To extend these observations, we asked whether CASM drives p62 recruitment in any other contexts, including drug treatment, agonist stimulation and entosis, a cancer-associated cell engulfment process ^37^. Notably, CASM-dependent p62 recruitment was clearly observed in response the lysosome stressor chloroquine, and agonists of TRPML1 (C8) and STING (diABZi) (Fig. S3A, B). However, upon closer examination, we noticed that the GFP-LC3A structures induced by different stimuli exhibited somewhat variable levels of p62 colocalisation. While LLOMe treatment resulted in strong co-localisation, monensin induced a more mixed phenotype, with both p62-positive and p62-negative GFP-LC3A structures apparent (Fig. S3C, D). Furthermore, in the context of entosis, we observed no detectable p62 recruitment to CASM-modified entotic vacuoles, decorated with GFP-LC3A (Fig. S3E). Together, these findings suggest that CASM drives p62 recruitment in response to lysosome stress, but that neither CASM activation, nor ATG8ylation of endosomal membranes, is sufficient to engage p62 alone, indicating that additional, stimulus-specific factors must also contribute to this process.

### CASM regulates ubiquitination of damaged lysosomes

Recruitment of selective autophagy receptors (SARs) to damaged lysosomes is generally thought to be driven by their binding to ubiquitinated targets at the lysosomal membrane ^13^. As such, we examined the impact of CASM on the ubiquitin response to LLOMe. In ATG13 KO cells, LLOMe and monensin both induced clear ubiquitin puncta, that colocalised with GFP-LC3A (Fig. 2A, B) and include K63-linked chains (Fig. S4A). Strikingly, inhibition of ATG8 lipidation, using ATG7i, significantly reduced ubiquitin recruitment. The reduction in lysosomal ubiquitination with the ATG7 inhibitor was also seen in wild type MCF10A cells not over-expressing GFP-LC3A (Fig. S4B), and a similar reduction was observed in ATG5 KO cells, also deficient in ATG8 lipidation (Fig. S4C). Importantly, we note this was a reduction and not a complete inhibition of ubiquitination. These data suggest that the magnitude of ubiquitin recruitment, which is typically considered an upstream event following lysosome damage, can be regulated by ATG8-lipidation.

**Figure 2.**
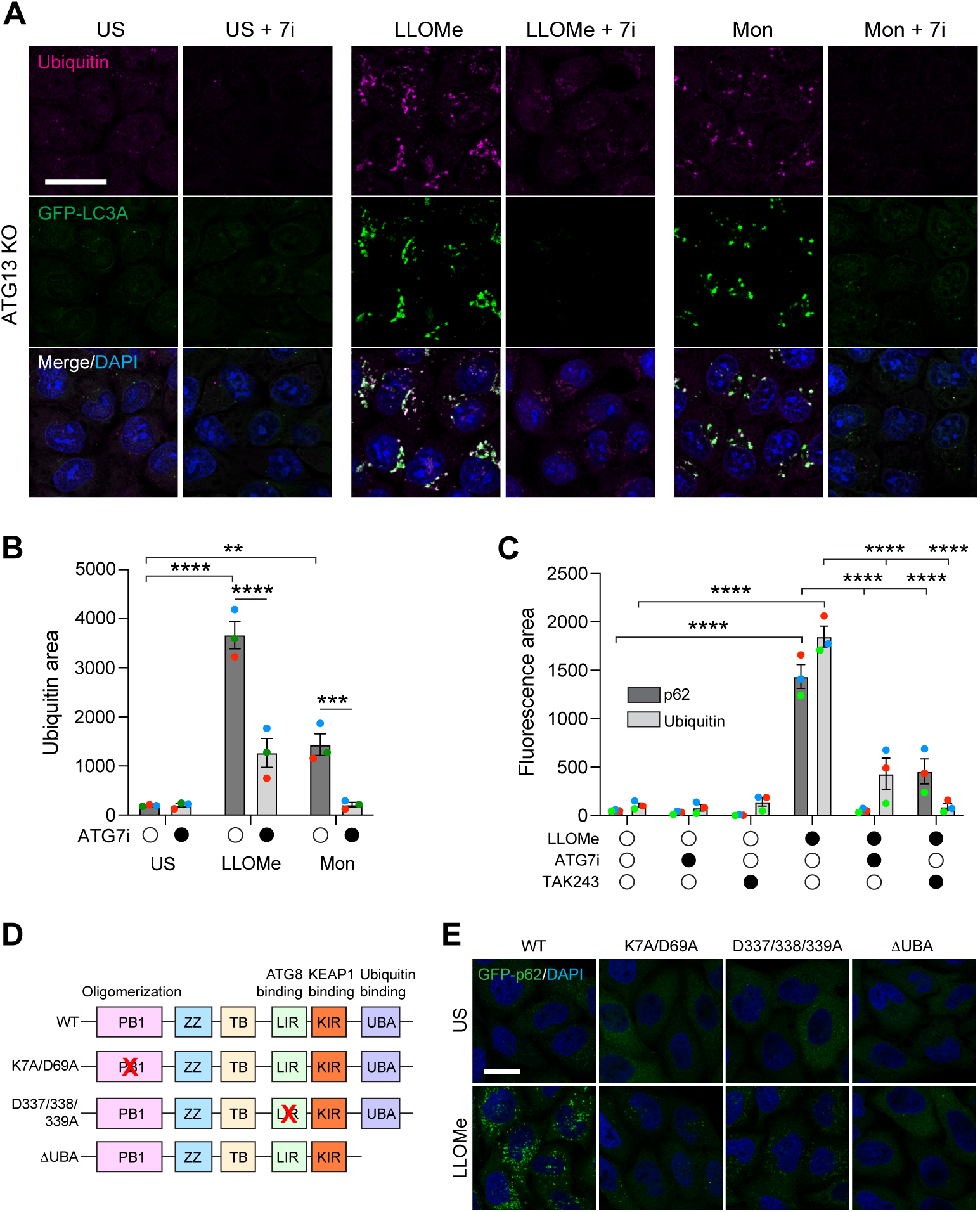
CASM regulates ubiquitination during lysosome damage, which is required for p62 recruitment. (A) Confocal images of ubiquitin, DAPI and GFP-LC3A in ATG13 KO MCF10A cells, unstimulated (US) or treated with LLOMe (250 μM, 30mins) or monensin (Mon, 100 μM, 45 min) +/- pretreatment with ATG7 inhibitor (7i, 10 μM, 2 hr). Scale bar: 25 μm. (B) Analysis of ubiquitin puncta area from images in (A). Data represent means ± SEM from 3 independent experiments. **** p<0.0001, *** p<0.0007, ** p 0.0018, 2 way anova with sidaks multiple comparison. (C) Analysis of p62 and ubiquitin puncta area from ATG13 KO MCF10A cells treated with LLOMe (250 μM, 30mins) +/- pretreatment with ATG7 inhibitor (7i, 10 μM, 2 hr) or E1 ligase inhibitor (TAK243, 5 μM, 30 min). Data represent means ± SEM from 3 independent experiments. **** p<0.0001, 2 way anova with sidaks multiple comparison. (D) Diagram of p62 structure showing relevant domains and key mutations. (E) Confocal images of p62 KO Hela cells re-expressing the indicated GFP-p62 constructs, unstimulated (US) or treated +/- LLOMe (250 μM, 30 min). Scale bar: 20 μm.

To further dissect the contributions of ATG8 lipidation and ubiquitination to p62 recruitment, we employed inhibitors of both ATG7 (ATG7i) and the ubiquitin-activating enzyme (E1) (TAK-243). Inhibition of either pathway led to a marked reduction in both ubiquitin and p62 puncta formation (Fig. 2C), consistent with a model in which ATG8-lipidation is required for efficient ubiquitin recruitment, which is in turn required for p62 engagement.

To further substantiate this model, we utilised p62 KO HeLa cells stably reconstituted with either wild-type GFP-p62 or mutants lacking key functional domains, including the PB1 oligomerisation domain (K7A/D69A), the LC3-interacting region (LIR, D337/338/339A), or the ubiquitin-associated (UBA) domain (ΔUBA, 1-388). While LLOMe treatment robustly induced GFP-p62 puncta in cells expressing wild-type p62, puncta formation was diminished across all mutant constructs (Fig. 2D, E). Some GFP-p62 puncta were detected in the LIR mutant, but these were fewer and less intense, which is consistent with previous findings that p62 LIR mutants still localise to autophagosomes ^38^. These data indicate that p62 recruitment to damaged lysosomes depends upon its binding to both ubiquitin and ATG8 (and the ability to oligomerise).

Collectively, these findings suggest that CASM plays an unexpected upstream role in promoting lysosomal ubiquitination following damage, and that efficient p62 recruitment requires both ATG8 lipidation and ubiquitin binding.

### CASM regulates the p62/KEAP1 axis in response to lysosome damage

To investigate the functional significance of CASM-mediated p62 recruitment, we examined phosphorylation at the regulatory S349 site in p62. Confocal microscopy revealed that LLOMe treatment increased pS349-p62 puncta in both wild-type and ATG13 KO cells, but not in ATG5 KO cells, which are deficient in ATG8 lipidation (Fig. 3A). Consistent with this, immunoblot analysis confirmed that S349 phosphorylation is enhanced upon LLOMe treatment, in wild-type and ATG13 KO cells, in a manner sensitive to ATG7 inhibition; no increase was observed in ATG5 KO cells (Fig. 3B). Together, these data suggest that p62 phosphorylation at S349 is dependent on CASM.

**Figure 3.**
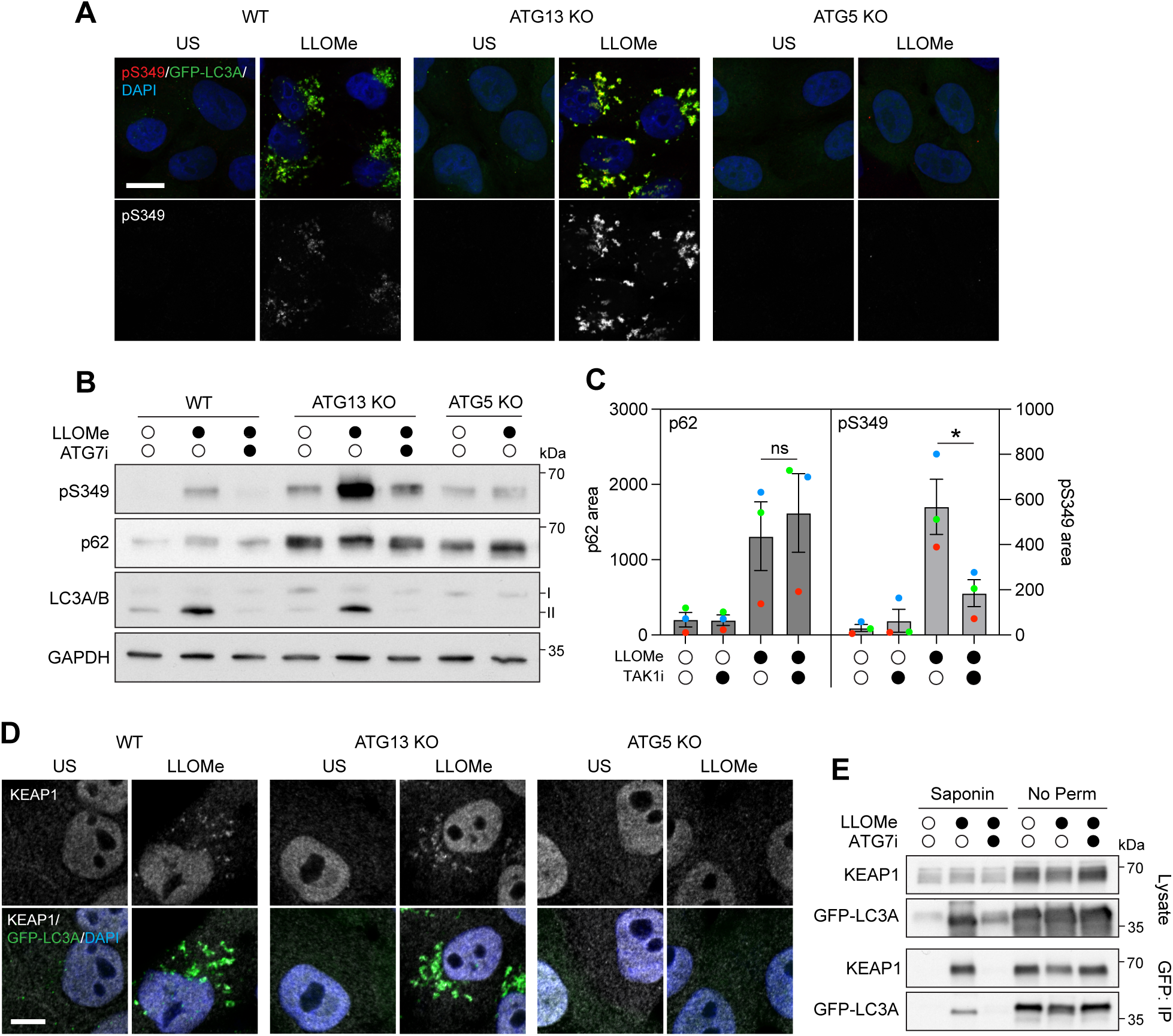
CASM regulates p62 phosphorylation and KEAP1 binding. (A) Confocal images of pS349 p62, DAPI and GFP-LC3A in wild type, ATG13 KO and ATG5 KO MCF10A cells, unstimulated (US) or treated +/- LLOMe (250 μM, 30 mins). Scale bar: 20 μm. (B) Immunoblot analysis of pS349, p62, LC3A/B and GAPDH in wild type, ATG13 KO and ATG5 KO MCF10A cells treated with LLOMe (250 μM, 30 min) +/- ATG7i pretreatment (10 μM, 2 hr). Scale bar: 5 μm. (C) Analysis of p62 and pS349 puncta area from ATG13 KO MCF10A cells treated with LLOMe (250 μM, 30mins) +/- pretreatment with TAK1 inhibitor (TAK1i, 10 μM, 30 min). Data represent means ± SEM from 3 independent experiments. * p 0.0025, 1 way anova with Tukey’s multiple comparison. (D) Confocal images of KEAP1, DAPI and GFP-LC3A in wild type, ATG13 KO and ATG5 KO MCF10A cells treated +/- LLOMe (250 μM, 30 mins). Scale bar: 5 μm. (E) Immunoblot analysis of KEAP1 and GFP-LC3A from total lysates and GFP-immunoprecipitation (GFP-IP) from ATG13 KO MCF10A cells treated with LLOMe (250 μM, 30 min) +/- ATG7i pretreatment (10 μM, 2 hr). Where indicated, cells were briefly permeabilised with saponin prior to lysis.

Several kinases can phosphorylate p62 at S349 ^39,40^, including TAK1 ^41,42^, which is also known to localise to damaged lysosomes in complex with TAB1/2, via ubiquitin binding ^43^. As such, we assessed TAK1 as a candidate kinase in this context. While inhibition of TAK1 did not affect overall p62 recruitment in LLOMe treated cells, it significantly reduced its S349 phosphorylation, as determined by confocal microscopy analysis (Fig. 3C), consistent with TAK1-dependent phosphorylation of p62, occurring post-recruitment.

Phosphorylation of p62 at S349 is known to enhance its affinity for KEAP1, a transcriptional regulator which functions during cellular stress; this modification promotes KEAP1 sequestration during selective autophagy ^39,44^. Analogously, we observed recruitment of KEAP1 to CASM-induced, GFP-LC3A-positive structures following lysosomal damage, dependent on ATG8 lipidation, but not ATG13 (Fig. 3D). Co-immunoprecipitation experiments further revealed the formation of an inducible complex between GFP-LC3A and KEAP1 upon lysosomal damage (Fig. 3E). In this experiment, saponin pre-treatment removes cytosolic, unlipidated ATG8s allowing for the specific analysis of proteins interacting with the lipidated ATG8 pool. This membrane-associated complex forms in ATG13 KO cells, dependent on ATG7, and thus bears the hallmarks of CASM. As KEAP1 is not thought to directly interact with LC3A ^44,45^, this association is likely mediated via S349-phosphorylated p62, bridging KEAP1 to LC3A.

Taken together, these data indicate that CASM-mediated p62 recruitment is coupled to its phosphorylation at S349, at least in part via TAK1, which promotes KEAP1 binding and sequestration at damaged lysosomes.

### CASM regulates the NRF2 antioxidant response to lysosome damage

Phosphorylation of p62 at S349, and its subsequent interaction with KEAP1, represents a well-established, non-canonical pathway for activation of NRF2, a master transcription factor that orchestrates the antioxidant response to cellular stress and damage ^39,44^. Given our new findings that lysosomal damage promotes CASM-dependent activation of the p62-KEAP1 axis, we examined the possible impact on NRF2 responses. Strikingly, confocal microscopy revealed that LLOMe treatment induced nuclear localisation of NRF2, in both wild-type and ATG13 KO cells, which was abrogated by ATG7 inhibition (Fig. 4A, B), consistent with its CASM-dependent nuclear translocation.

**Figure 4.**
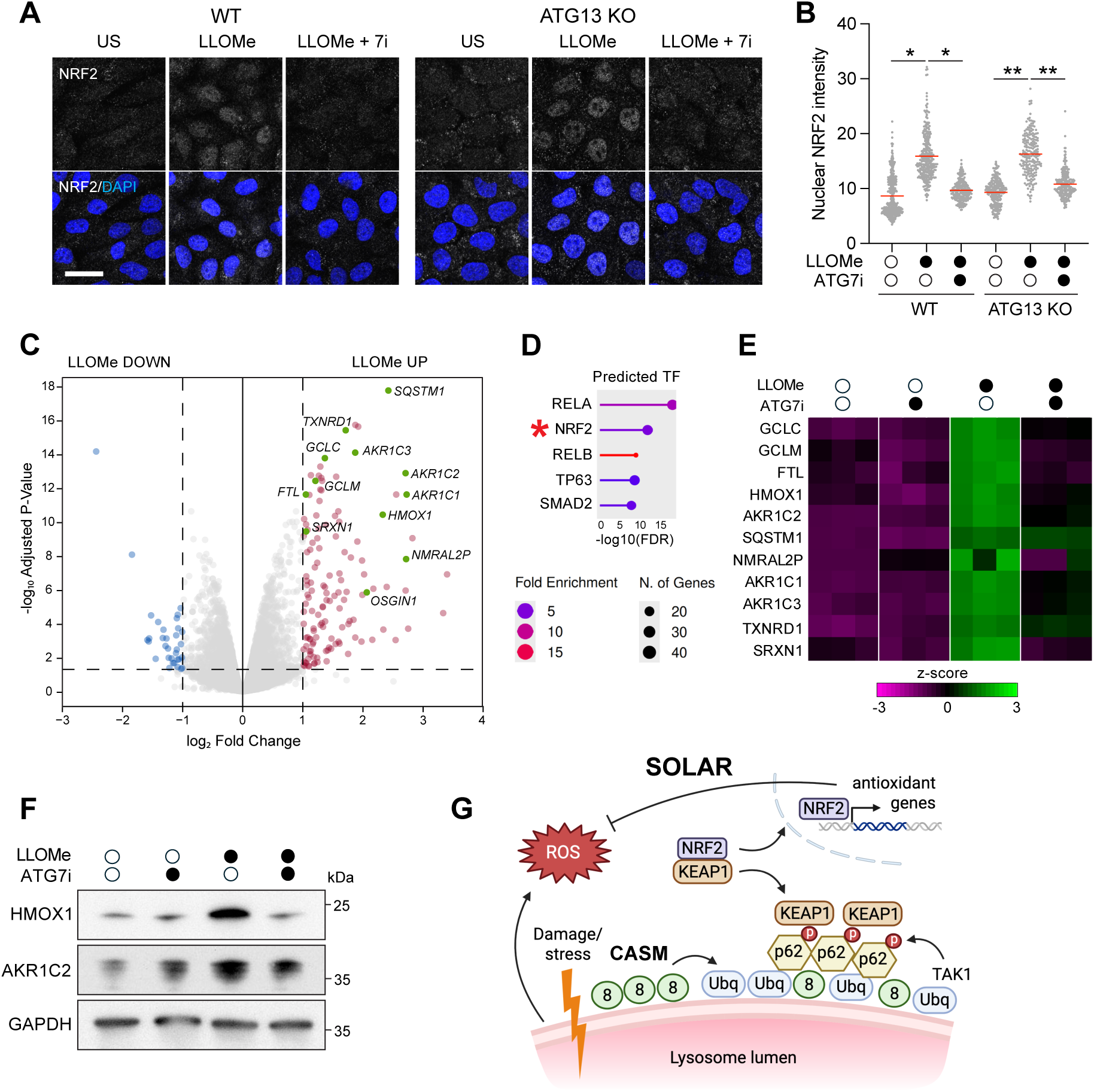
CASM regulates NRF2 activation and antioxidant transcriptional response. (A) Confocal images of NRF2 and DAPI in wild type and ATG13 KO MCF10A cells, unstimulated (US) or treated +/- LLOMe (250 μM, 6 hr) +/- ATG7i pretreatment (10 μM, 2 hr). Scale bar: 30 μm. (B) Analysis of nuclear NRF2 intensity in cells treated as in (A). Data represent individual nuclei and mean from 3 independent experiments. ** p<0.009, * p<0.03, mixed effect analysis with Tukey’s multiple comparison test. (C) The mean Log2 fold change (FC, LLOMe 250 μM, 6 h/control) and −Log10 *P* value of the transcriptome in ATG13 KO MCF10A cells are indicated on the x and y axes, respectively. Genes upregulated or downregulated by LLOMe treatment are labeled in red and blue, respectively. NRF2 target genes are annotated in green. (D) The bubble plot illustrates Transcription Factor Enrichment Analysis for transcription factors responsible for the induction of genes upregulated in cells treated with LLOMe (top five transcription factors). The *x* axis represents the -Log10(FDR) and the color and size of the bubbles indicate the fold enrichment and number of genes respectively. (E) Heatmap displaying the z-score expression pattern of a curated set of NRF2 target genes from cells treated +/- LLOMe (250 μM, 6 hr), +/- ATG7i pretreatment (10 μM, 2 hr). (F) Immunoblot analysis of HMOX1, AKR1C2 and GAPDH in cells treated as in (E). (G) Cartoon depicting the upstream role of CASM (8 = ATG8) in regulating ubiquitin (Ubq) and p62 recruitment upon lysosome damage, driving SOLAR-mediated KEAP1 sequestration and NRF2 activation. Created with Biorender.com.

To characterise the resultant transcriptional response to lysosomal damage, and assess its dependence on CASM, we performed RNA-seq analysis, using ATG13 KO cells to exclude macroautophagy and lysophagy, and ATG7i to modulate ATG8-lipidation. Numerous genes were significantly upregulated in response to lysosome damage (Fig. 4C), and transcription factor enrichment analysis identified NRF2 as a key regulator (Fig. 4D). Focusing on a curated set of NRF2 target genes, z-score analysis demonstrated that LLOMe increased their expression, in a manner dependent on ATG7 (Fig. 4E). Building on these findings, immunoblot analysis of representative NRF2 target genes, HMOX1 (heme oxygenase 1) and AKR1C2 (aldo-keto reductase), confirmed that LLOMe increased expression at the protein level, in the absence of ATG13, and dependent on ATG7 (Fig. 4F), and thus consistent with CASM dependency. HMOX1 and AKR1C2 represent classic stress-response genes involved in antioxidant, detoxification and cytoprotective pathways, suggesting that CASM activates a transcriptional program to mitigate lysosome stress and respond to damage.

Together, these new findings establish a novel, and somewhat surprising, upstream role for the CASM pathway following lysosome damage, mediating ubiquitination, p62 recruitment and S349 phosphorylation, and ultimately driving the KEAP1/NRF2 transcriptional antioxidant response.

## Discussion

Membrane ATG8ylation represents a major cellular response to lysosome stress. In current models of lysophagy, ATG8 lipidation occurs downstream of lysosomal ubiquitination and selective autophagy receptor (SAR) recruitment ^13^. Here, we discover a novel upstream function, mediated by the direct, CASM-dependent ATG8ylation of damaged endolysosomal membranes. In this context, CASM-induced ATG8ylation is necessary for efficient lysosome ubiquitination and subsequent activation of a p62/KEAP1/NRF2 signalling axis, which drives transcription of antioxidant genes. To distinguish p62’s function here from its established role in selective autophagy, we refer to this lysosome stress-specific activity as SOLAR (SQSTM1/p62 oligomer-mediated lysosomal antioxidant response). CASM-mediated SOLAR orchestrates the transcription of various antioxidant and stress response genes, including HMOX1 and AKR1C2, thereby mounting a cytoprotective response. This model is summarised in (Fig. 4G).

Mechanistically, we have also gained new insights into the roles of SAR proteins during lysosome stress. In the context of lysophagy, OPTN, TAX1BP1, and NDP52 play key roles in clearance ^19^, but the relative contributions of other SARs remain incompletely resolved; the role of p62 appears more variable and may depend on cell type or context ^46,47^. Here, we propose that p62 fulfils an alternative, upstream function following lysosome damage, acting in the SOLAR pathway, downstream of CASM. We note that inhibition of CASM and p62 recruitment in KI/KO cells did not impair lysophagy-associated autophagosome formation (Fig. S1E), suggesting these events are mechanistically, and probably temporally, uncoupled. Interestingly, among other SARs, NBR1 is similarly recruited, albeit at a lower level, to damaged lysosomes in a CASM-dependent manner, but NDP52 is not, indicating selectivity, and possible division of functions among different SAR family members over the course of the lysosome damage response.

A key outstanding question raised by this study is how CASM regulates ubiquitination of damaged lysosomes. Importantly, CASM modulates the extent of lysosomal ubiquitination rather than being strictly required for it. We hypothesise that lipidated ATG8s may engage one or more LIR-domain containing E3-ubiquitin ligases, and therefore would specifically regulate the ubiquitination of their target substrates as opposed to total ubiquitination levels. The identity(s) of these ligases, including candidates such as NEDD4/L, SMURF1, TRIM5/13/16/21, Parkin, MARCH5, RNF185, and their lysosomal substrates, represent a compelling area for future research. Notably, p62 and NBR1 both bear a canonical C-terminal UBA ubiquitin-binding domain, while NDP52 uses a zinc-finger ubiquitin-binding module, which is generally lower affinity and more context-dependent ^48^; these differences may underpin the selective CASM-dependent recruitment of p62-like SARs.

In summary, we report a novel lysosome stress response pathway (SOLAR), in which CASM-dependent lysosome ubiquitination and p62 recruitment activate the non-canonical p62-KEAP1-NRF2 signalling axis, driving a transcriptional programme enriched for antioxidant response genes. Lysosomal damage is known to promote reactive oxygen species (ROS) production and oxidative stress ^49^, and as such, SOLAR may function to limit secondary cellular damage following lysosomal injury. Thus, alongside its established, acute membrane repair functions, through the recruitment of ESCRT and ATG2 machinery ^30–33^, CASM also coordinates adaptive stress signalling responses to promote cellular recovery from lysosomal stress. These findings broaden the functional scope of CASM and identify p62 as a new, and somewhat unexpected, downstream CASM effector. More broadly, these findings also add to emerging evidence that CASM orchestrates transcriptional responses to lysosomal stress, complementing the discovery of CASM-dependent TFEB activation, to support lysosome biogenesis ^34^. Together, these observations position CASM as a central lysosomal stress response pathway that integrates membrane repair, stress signalling, and multiple transcriptional programmes to maintain cellular homeostasis following lysosomal damage.

## Acknowledgements

We thank all members of the Florey Lab for their input, and the Babraham Institute Imaging and Bioinformatics facilities for excellent technical support. We thank Leon Murphy (Casma Therapeutics) for kindly providing the ATG7 inhibitor and TRPML1 agonist C8, and Viktor Korolchuk (Newcastle University) for generously sharing p62 KO Hela cells. This work was supported by a BBSRC (Biotechnology and Biological Sciences Research Council) institutional programme grant (BB/Y0069251/1) and by a BBSRC CTP Studentship (BB/X511535/1). Use of the Institute Bioinformatics and Imaging facilities was supported by the Babraham Institute’s UKRI-BBSRC Core Capability Grant

## Declarations of interest

RS is a founder, CEO, CSO and shareholder in Stemnovate Limited. All other authors declare no competing interests.

## Methods

### Reagents

Antibodies used were rabbit polyclonal anti-LC3A/B (Cat#4108, WB 1:1,000, IF 1:100; Cell Signalling), mouse monoclonal anti-GAPDH (Cat#ab8245, WB 1:1,000; Abcam), mouse anti-GFP (Cat#1181446000, WB 1:1,000; Roche), mouse anti-WIPI2 (Cat#MCA5780GA, IF 1:100; BioRad), mouse monoclonal anti-p62 (Cat#610833, WB 1:1000, IF 1:100; BD Biosciences), rabbit polyclonal anti-pS349 p62 (Cat#16177, WB 1:1000, IF 1:100; Cell Signalling), mouse monoclonal anti-ubiquitin (Cat#SMC-214, IF 1:100; StressMarq), rabbit polyclonal anti-KEAP1 (Cat#10503-2-AP, WB 1:2000, IF 1:100; ProteinTech), rabbit polyclonal anti-NRF2 (Cat#12721, WB 1:1000, IF 1:200; Cell Signalling), rabbit polyclonal anti-AKR1C2 (Cat#13035S, WB 1:1000; Cell Signalling), rabbit polyclonal anti-HMOX1 (Cat#10701-1-AP, WB 1:5000; ProteinTech), rabbit polyclonal anti-TECPR1 (Cat#8097, WB 1:500; Cell Signalling), rabbit polyclonal anti-ATG16L1 (Cat#8089, WB 1:1,000; Cell Signalling). Alexa Fluor 488 polyclonal goat anti-rabbit IgG (Cat#A-11034, IF 1:500; Thermo Fisher Scientific), Alexa Fluor 568 polyclonal goat anti-mouse IgG (Cat#A-11004, IF 1:500; Thermo Fisher Scientific), Alexa Fluor 568 polyclonal goat anti-rabbit IgG (Cat#A-11011, IF 1:500; Thermo Fisher Scientific), HRP-conjugated anti-rabbit IgG (Cat#7074, WB 1:2,000; Cell Signalling), and HRP-conjugated anti-mouse IgG (Cat#7076, WB 1:2,000; Cell Signaling Technology).

Reagents and chemicals used were LLOMe (L7393; Sigma-Aldrich), BafA1 (1334; Tocris), PP242 (4257; Tocris), Monensin (M5273; Sigma-Aldrich), DAPI (D9542; Sigma-Aldrich), IN-1 (17392; Caymen Chemical), diABZi (tlrl-diABZi-2; Invivogen SAS), chloroquine (C6628; Sigma-Aldrich), TAK243 (HY-100487; Cambridge Bioscience), TAK1i (CAY-24161; Cambridge Bioscience), saponin (84510; Sigma-Aldrich), puromycin (P8833; Sigma-Aldrich). TRPML1 agonist C8 and ATG7 inhibitor ATG7-IN-3 were kindly provided by Casma Therapeutics. GFP-Trap (gtma-20), and control magnetic agarose beads (bmab-20) were obtained from Chromotek.

### Cell culture

WT, ATG13 KO and ATG5 KO (Cat#CRL-10317, RRID:CVCL_0598; ATCC) expressing GFP-LC3A (human) were prepared as described previously ^50–52^ and cultured in DMEM/F12 (11320074; Gibco) containing 5% horse serum (16050-122; Thermo Fisher Scientific), EGF (20 ng/ml, AF-100-15; Peprotech), hydrocortisone (0.5 mg/ml H0888; Sigma-Aldrich), cholera toxin (100 ng/ml, C8052; Sigma-Aldrich), insulin (10 μg/ml, I9278; Sigma-Aldrich), and penicillin/streptomycin (100 U/ml 15140-122; Gibco) at 37°C, 5% CO2.

Wild type and p62 KO Hela cells were maintained in DMEM (41966-029; Gibco) supplemented with 10% FBS (F9665; Sigma-Aldrich) and penicillin/streptomycin (100 U/ml, 100 μg/ml 15140-122; Gibco) at 37°C, 5% CO2.

### Genome editing with CRISPR Cas9

ATG16L1 K490A Knockin MCF10A cells were generated using the Lipofectamine CRISPRmax/Cas9 system (ThermoFisher Scientific), with single strand guide RNAs, and donor DNA sequence designed using TrueDesign genome Editor. Briefly, cells in antibiotic free media were transfected with guide (7.5 pmoles) (guide: GTTGGGAAAGATTACTGCCC), donor DNA (10 pmol) (donor: CAGAGAGCATAGTTCGAGAGATGGAGCTGTTGGGAGCCATTACGGCCCTTGACTTAAACCCAGAAAGGACTGAGCTCCTG), Cas9 protein (1.25 ug TrueCut Cas9 Protein v2 A36497) and Lipofectamine CRISPRMAX Reagent and Cas9 Plus Reagent (CMAX00003; ThermoFisher) as per manufacture instructions, in the presence of HDR enhancer V2 (IDT; 10007910, 1 uM final). Media was replaced 24 hr post-transfection to remove HDR. Single cell clones, sorted by FACS, were expanded and K490A knockin confirmed by sequence analysis using PCR primers shown in (Forward primer: 5’TGGGAATGACTGTCTTAGGGTCTGT ; Reverse primer: 5’CCTGGAAACATAGGTTGGGCACT).

TECPR1 was knocked out in MCF10A K490A cells using a TECPR1 guide targeting exon 3 (guide: GCGTGTACACTCTCCCGAAG), to generate KI/KO cells using method described above, with the absence of donor DNA and HDR enhancer in the transfection. Sequence frameshift and knockout of TECPR1 in flow sorted single cell derived clones was confirmed by sequencing (DNS) (Forward primer: 5’GTCTCCCAGGTTTCACATGAC; Reverse primer: 5’CCACGCCTGGCAGCAAC) and Western blot analysis.

For p62 KO, HeLa cells were seeded at 2.5x10^5^ cells in a 6-well plate-well and transfected the following day with sgRNA constructs targeting p62 exon 3 (gift from Lazarou lab), hCas9 plasmid (Addgene #41815), and pEGFP-C1 (Clonetech) as a transfection marker using lipofectamine 3000 standard protocol. After 48 hours, GFP-positive cells were isolated by FACS and single cells were distributed into 96-well plates for clonal expansion. Successfully expanded clones were transferred to 24-well plates for further growth. Knockout efficiency was assessed by Western blot analysis.

### GFP-p62 cloning

pMXs-puro GFP-p62 (mouse) constructs were a gift from Noboru Mizushima and purchased from addgene (wild type #38277; K7A/D69A #38281; D337/338/339A #38280). To generate GFP-p62-ΔUBA (1-388), PCR was used to introduce a stop codon at position 388, along with 5′ EcoRI and 3′ KpnI restriction sites, into wild-type GFP-p62 (Forward primer: 5’CGATGAATTGAATTCCATGGCGTCG; Reverse primer: 5’GCTGCGGTACCTCAATGTGGGTATAGGGCAGCT). The PCR product was then cloned into pMXs-puro-GFP-p62 using EcoRI/KpnI restriction digest.

### Retrovirus generation and infection

pMXs-puro-GFP-p62 constructs were transiently transfected into HEK293T cells along with envelope and packaging constructs using Lipofectamine 2000 (Invitrogen) following the manufacturer’s guidelines. Viral supernatant was collected over 2 days. For infection, Hela cells were seeded in a six-well plate at 5 × 10^4^ per well. The next day, 1 ml viral supernatant was added with 10 µg/ml polybrene for 24 hr followed by a media change. Cells were then selected with puromycin (2 µg/ml). Cells were then flow sorted to enrich cells with similar GFP expression levels.

### Immunofluorescence

Cells were seeded in 12-well plates containing coverslips and incubated at 37°C, 5% CO2 for 24 hr. Following treatments, cells were washed twice with ice-cold PBS and then incubated with 100% methanol at −20°C for 10 min. Alternatively, cells were fixed with 4% formaldehyde at RM temp for 10 mins followed by permeabilization with 0.2% TritonX100 in PBS for 10 min. The cells were then washed twice with PBS and blocked with 5% BSA (A7906; Sigma-Aldrich) in PBS for 1 hr at room temperature. The cells were incubated overnight at 4°C with the primary antibodies in blocking buffer and then washed ×3 in cold PBS. Fluorescent secondary antibodies were used at a 1:500 dilution in blocking buffer and were incubated with the cells for 1 hr at room temperature. The cells were washed ×3 in cold PBS prior to being incubated with DAPI for 10 min and then mounted onto microscope slides with ProLong Gold anti-fade reagent (P36930; Invitrogen). Image acquisition was performed with the Confocal Zeiss LSM 780 microscope (Carl Zeiss Ltd) equipped with a 40x oil immersion 1.40 numerical aperture (NA) objective using Zen software (Carl Zeiss Ltd).

### Image analysis

Fluorescence area quantification was performed using ImageJ (RRID:SCR_003070). Images were batch processed using a custom macro. Briefly, .czi files were imported using the Bio-Formats importer as hyperstacks. Images were then thresholded using the “Threshold” tool at a fixed intensity (number selected as appropriate for that data set) to ensure consistency across all samples. A binary selection was generated from the thresholded signal, and the total fluorescent area was measured using the “Measure” function in Fiji. The total fluorescence area values for each image were exported to a .csv file using the custom macro. Cell number per field of view was then counted manually. To account for differences in cell density, the total fluorescence area per image was divided by the corresponding cell number, yielding fluorescence area per cell.

For NRF2 nuclear density analysis, individual nuclei regions were selected by thresholding in the DAPI channel using ImageJ. NRF2 signal intensity was then measured within nuclei regions.

### Cell lysis and GFP-TRAP immunoprecipitation

For routine cell lysis, cells were seeded at 2 × 10⁵ cells per well in a 6-well plate and culured for 48 h. Cells were washed with ice-cold PBS and scraped into 80 µl of RIPA buffer (150 mM NaCl, 50 mM Tris–HCl (pH 7.4), 1 mM EDTA, 1% Triton X-100, 0.1% SDS, 0.5% sodium deoxycholate) supplemented with phosphatase inhibitors (1×, P0044; Sigma-Aldrich) and protease inhibitors (1×, P8340; Sigma-Aldrich). Lysates were incubated on ice for 10 min before centrifugation at 10,000 × rpm for 10 min. The resulting supernatant was combined with 5x sample buffer and heated at 95°C for 5 min.

For GFP-TRAP, cells from a 100 mm dish were pelleted at 200 x g for 5min and washed once in ice-cold PBS. Cells to be permeabilised were resuspended in 1.8 ml of ice-cold permeabilisation buffer (PBS supplemented with 0.5% saponin) in a pre-chilled 2 ml microcentrifuge tube and incubated on ice for 5 min. Cells were then pelleted and washed twice in ice-cold PBS by centrifugation at 200 × g for 5 min at 4°C.

Cells were lysed in 500 µl of ice-cold lysis buffer [50mM Tris HCl (pH 7.5) 150mM NaCl, 1% Triton X-100] supplemented with phosphatase inhibitors (1×, P0044; Sigma-Aldrich) and protease inhibitors (1×, P8340; Sigma-Aldrich) for 10 min on ice, and lysates underwent centrifugation at 13,000 rpm for 10min at 4°C. A 20 µl aliquot of the resulting supernatant was reserved as the total cell lysate input by addition to 20 µl of 2× LDS sample buffer, heated at 95°C for 5 min, and stored at −20°C. The remaining supernatant was transferred to pre-equilibrated control beads and rotated at 4°C for 30 min. Beads were immobilised using a magnetic rack and the supernatant was transferred to pre-equilibrated GFP-trap beads. The suspension was rotated at 4°C for 1 h to allow immunoprecipitation.

Beads were then washed three times in 700 µl of ice-cold lysis buffer, resuspended in 30 µl of 2× LDS sample buffer, heated at 95°C for 5 min, and stored at −20°C.

### Western blotting

Western blotting was carried out as previously described (Hooper et al., 2022). In brief, cell lysates were separated by SDS-PAGE and transferred onto PVDF membrane (Immobilon-P; Millipore). Membranes were blocked with 5% BSA/TBS-T for 1 h at room temperature, followed by overnight incubation with primary antibody at 4°C. After 3 x 10min washes in TBS-T, membranes were incubated with HRP-conjugated secondary antibodies for 45 min at room temperature. Membranes were then washed a further 3 x 10 min in TBS-T before development with ECL (RPN2106; GE). Blots were developed using an MI-5 X-Ray film processor (Medical Index). For TECPR1, lysates were run on 4-12% gradient SDS-PAGE gels (BioRad, 456-1085), transferred to PVDF membrane (Immobilon-P; Millipore) overnight at 21V, blocked with 5% Milk/TBS-T for 1 h, room temp, and then incubated with primary antibody at 4°C overnight in 5%BSA/TBS-T.

### mRNA-sequencing, bioinformatics, and data visualization

10^5^ cells in 50 ul Zymo DNA/RNA Shield (R110050; Cambridge Bioscience) were submitted to Plasmidsaurus. RNA sequencing was performed by Plasmidsaurus using a 3′-end counting approach. Briefly, poly(A)+ RNA was captured using oligo-dT priming, followed by reverse transcription incorporating unique molecular identifiers (UMIs), second-strand synthesis, and tagmentation-based library preparation. Libraries were PCR-amplified with dual indices and sequenced on an Illumina platform to generate single-end ∼90 bp reads. Sequencing yielded approximately 20 million raw reads per sample, with ∼10 million deduplicated reads retained following UMI-based collapsing. Raw reads were processed using a standard pipeline including quality filtering and adapter trimming (fastp), alignment to the reference genome using STAR, and UMI-based deduplication. Gene-level counts were generated from deduplicated alignments for downstream analysis. Transcription factor enrichment analysis was performed using ShinyGo using the ChEA 2022 dataset. Z-score analysis was performed using SeqMonk.

### Statistics

Statistical tests were performed using GraphPad, as indicated in figure legends.

