## Supplemental Figures for "CASM regulates p62/KEAP1/NRF2 antioxidant responses to lysosome damage"

Figure S1.
Figure S2.
Figure S3.
Figure S4.

### Figure S1

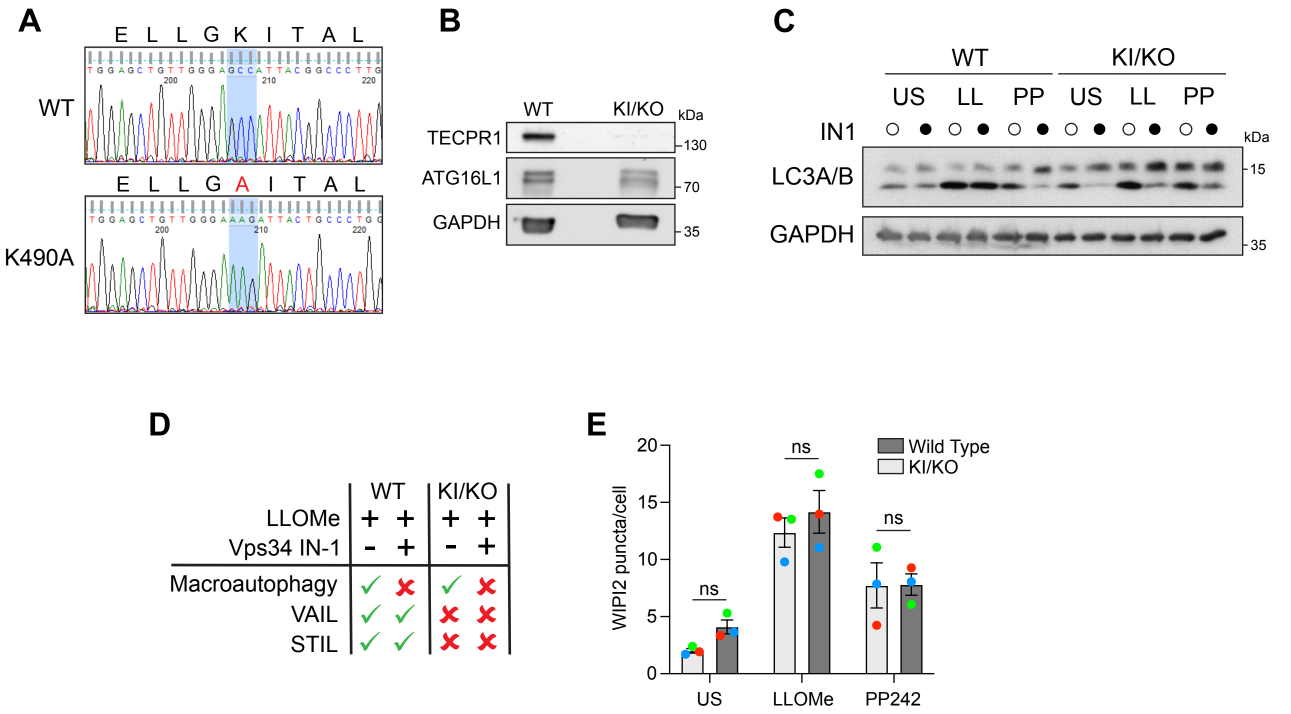
A
WT
K490A
E L L G K I T A L
E L L G A I T A L
B
WT
KI/KO
kDa
TECPR1
ATG16L1
GAPDH
130
70
35
C
WT
KI/KO
US LL PP US LL PP
IN1
LC3A/B
GAPDH
kDa
15
35
D
WT
KI/KO
LLOMe
Vps34 IN-1
Macroautophagy
VAIL
STIL
E
WIPI2 puncta/cell
ns
Wild Type
KI/KO
US
LLOMe
PP242

**Figure S1. Engineering of a CASM-deficient KI/KO MCF10A cell line (ATG16L1 K490A KI/TECPR1 KO)**

(A) Sequence alignment of WT and KI/KO MCF10A cells, demonstrating the knock-in of the K490A mutation.

(B) Immunoblot analysis of TECPR1, ATG16L1 and GAPDH in WT and KI/KO cells.

(C) Immunoblot analysis of LC3A/B and GAPDH in WT and KI/KO cells, treated with LLOMe (250 μM, 30 mins) or mTOR inhibitor (PP242, 1 μM, 1 hr), +/- pretreatment with Vps34 inhibitor (IN1, 5 μM, 30 min).

(D) Table indicating which ATG8 lipidation pathway is activated or inhibited under conditions used in (C).

(E) Analysis of autophagosome formation via quantification of WIPI2-positive puncta formation in WT and KI/KO cells, treated with LLOMe (250 μM, 30 mins) or mTOR inhibitor (PP242, 1 μM, 1 hr).

### Figure S2

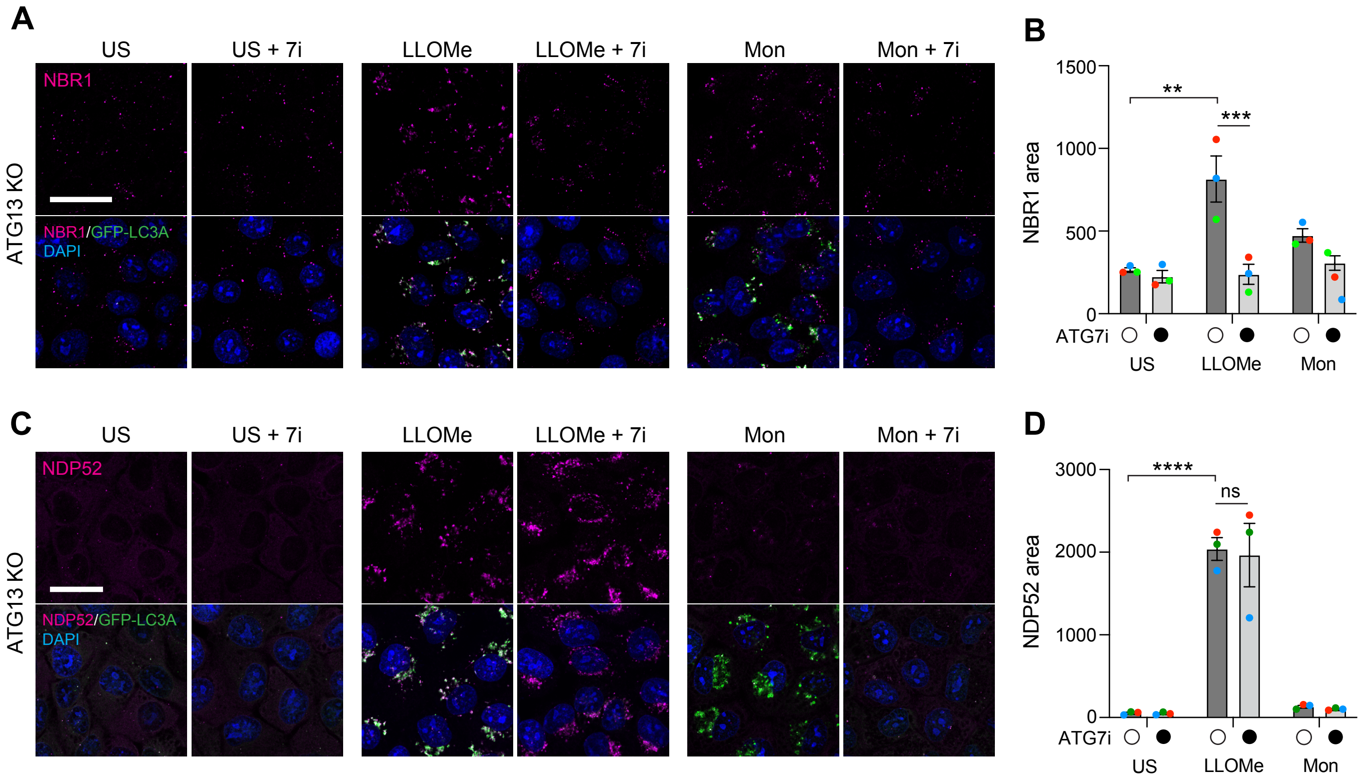
A
US
US + 7i
LLOMe
LLOMe + 7i
Mon
Mon + 7i
NBR1
ATG13 KO
NBR1/GFP-LC3A
DAPI
B
NBR1 area
1500
1000
500
0
**
***
ATG7i
US
LLOMe
Mon
C
US
US + 7i
LLOMe
LLOMe + 7i
Mon
Mon + 7i
NDP52
ATG13 KO
NDP52/GFP-LC3A
DAPI
D
NDP52 area
3000
2000
1000
0
****
ns
ATG7i
US
LLOMe
Mon

**Figure S2. Differential regulation of NBR1 and NDP52 recruitment to damaged lysosomes.**

(A) Confocal images of NBR1, DAPI and GFP-LC3A in ATG13 KO MCF10A cells, unstimulated (US) or treated with LLOMe (250 μM, 30mins) or monensin (Mon, 100 μM, 45 min) +/- pretreatment with ATG7 inhibitor (7i, 10 μM, 2 hr). Scale bar: 20 μm.

(B) Analysis of NBR1 puncta area from images in (A). Data represent means ± SEM from 3 independent experiments. ** p<0.0016, *** p 0.0010, 2 way anova with sidaks multiple comparison.

(C) Confocal images of NDP52, DAPI and GFP-LC3A in ATG13 KO MCF10A cells, unstimulated (US) or treated with LLOMe (250 μM, 30mins) or monensin (Mon, 100 μM, 45 min) +/- pretreatment with ATG7 inhibitor (7i, 10 μM, 2 hr). Scale bar: 25 μm.

(D) Analysis of NDP52 puncta area from images in (C). Data represent means ± SEM from 3 independent experiments. **** p<0.0001, 2 way anova with sidaks multiple comparison.

### Figure S3

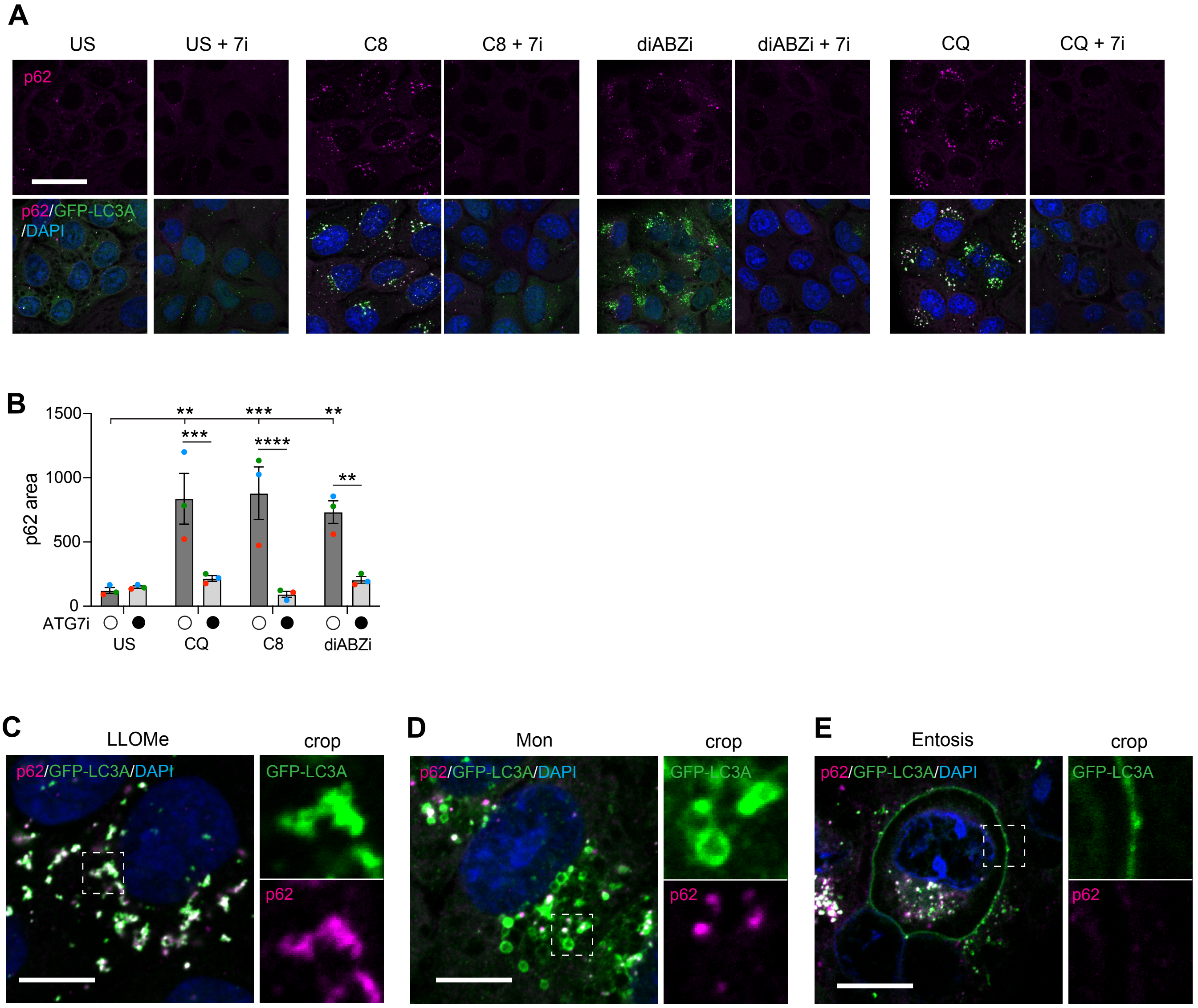
A
US
US + 7i
C8
C8 + 7i
diABZi
diABZi + 7i
CQ
CQ + 7i
p62
p62/GFP-LC3A
/DAPI
B
p62 area
1500
1000
500
0
**
***
**
***
****
**
ATG7i
US
CQ
C8
diABZi
C
LLOMe
crop
p62/GFP-LC3A/DAPI
GFP-LC3A
p62
D
Mon
crop
p62/GFP-LC3A/DAPI
GFP-LC3A
p62
E
Entosis
crop
p62/GFP-LC3A/DAPI
GFP-LC3A
p62

**Figure S3. A subset of CASM stimuli induce p62 puncta formation.**

(A) Confocal images of p62 and GFP-LC3A in ATG13 KO MCF10A cells treated with TRPML1 agonist C8 (2 μM, 30mins) or STING agonist diABZi (100 μM, 45 min) or chloroquine (CQ, 100 μM, 45 mins) +/- pretreatment with ATG7 inhibitor (7i, 10 uM, 2 hr). Scale bar: 30 μm.

(B) Analysis of p62 puncta area from images in (A). Data represent means ± SEM from 3 independent experiments. ** p<0.005, *** p 0.0008, **** p<0.0001, 2 way anova with sidaks multiple comparison.

(C-E) Confocal images of p62 and GFP-LC3A in MCF10A cells treated with (C) LLOMe or (D) monensin. (E) shows an entotic cell-in-cell structure with a GFP-LC3A positive entotic vacuole. Scale bar: 10 μm.

### Figure S4

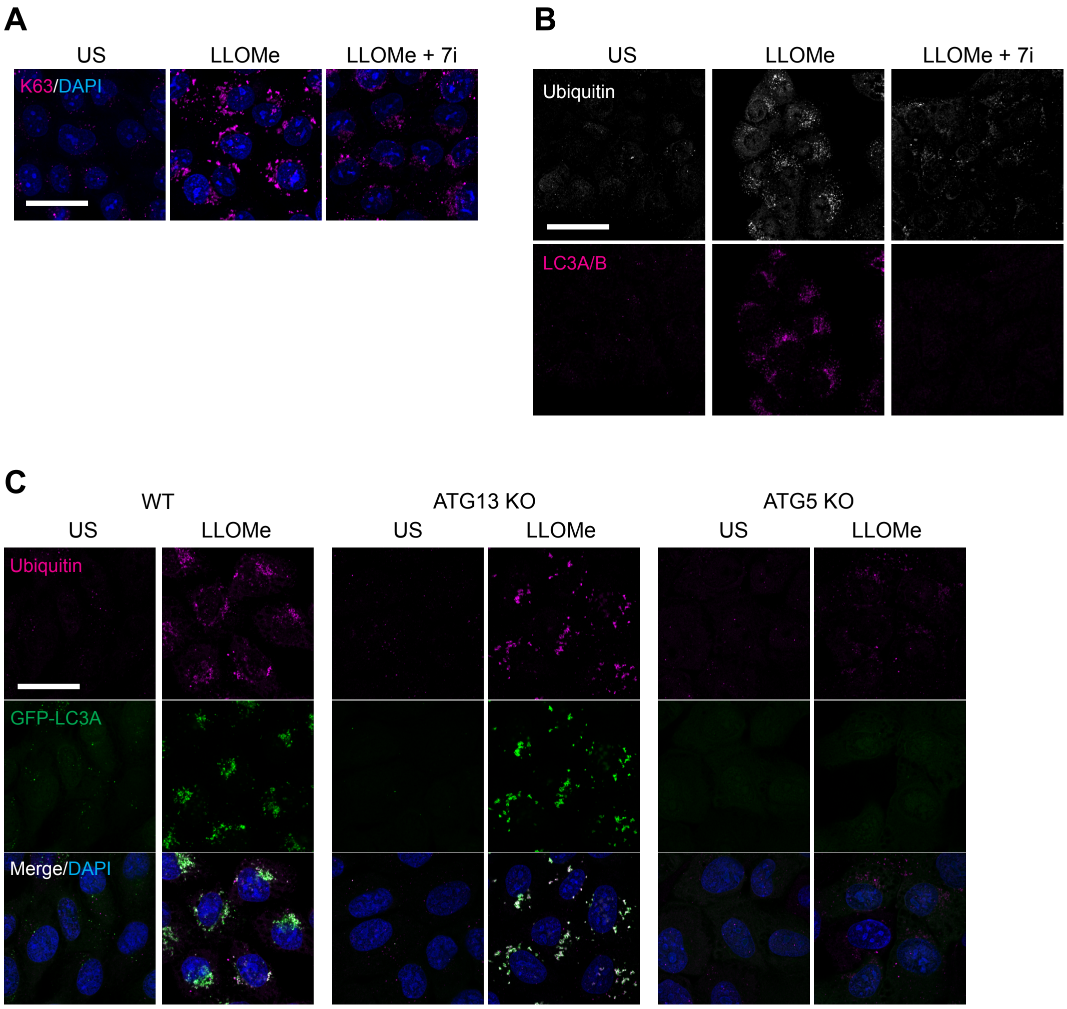
A
US
LLOMe
LLOMe + 7i
K63/DAPI
B
US
LLOMe
LLOMe + 7i
Ubiquitin
LC3A/B
C
WT
ATG13 KO
ATG5 KO
US
LLOMe
US
LLOMe
US
LLOMe
Ubiquitin
GFP-LC3A
Merge/DAPI

**Figure S4. Genetic inhibition of ATG8 lipidation inhibits ubiquitination of damaged lysosomes.**

(A) Confocal images of ubiquitin and GFP-LC3A in wild type, ATG13 KO and ATG5 KO MCF10A cells treated +/- LLOMe (250 μM, 30 mins). Scale bar: 25 μm.
(B) Confocal images of ubiquitin and endogenous LC3A/B in wild type cells treated with LLOMe (250 μM, 30 min) +/- ATG7i pretreatment (10 μM, 2 hr). Scale bar: 25 μm.
(C) Confocal images of K63-linked ubiquitin in MCF10A cells treated with LLOMe (250 μM, 30 min) +/- ATG7i pretreatment (10 μM, 2 hr). Scale bar: 25 μm.
